# A deletion at the X-linked *ARHGAP36* gene locus is associated with the orange coloration of tortoiseshell and calico cats

**DOI:** 10.1101/2024.11.19.624036

**Authors:** Hidehiro Toh, Wan Kin Au Yeung, Motoko Unoki, Yuki Matsumoto, Yuka Miki, Yumiko Matsumura, Yoshihiro Baba, Takashi Sado, Yasukazu Nakamura, Miho Matsuda, Hiroyuki Sasaki

**Affiliations:** Medical Institute of Bioregulation, Kyushu University, Fukuoka, Japan; National Institute of Genetics, Research Organization of Information and Systems, Mishima, Japan; Colledge of Liberal Arts, International Christian University, Mitaka, Japan; School of International Health, Graduate School of Medicine, The University of Tokyo, Tokyo, Japan; Data Science Center, Azabu University, Sagamihara, Japan; Research and Development Section, Anicom Specialty Medical Institute Inc., Yokohama, Japan; School of Medicine, Nagoya University, Nagoya, Japan; Inakazu Dog & Cat Kashii Hospital, Fukuoka, Japan; Graduate School of Agriculture, and Agricultural Technology and Innovation Research Institute, Kindai University, Nara, Japan; Faculty of Dental Science, Kyushu University, Fukuoka, Japan

## Abstract

The X-linked *orange* (*O*) locus in domestic cats controls an unknown molecular mechanism that causes the suppression of black-brownish pigmentation in favor of orange coloration. The alternating black-brownish and orange patches seen in tortoiseshell and calico cats are considered as classic examples of the phenotypic expression of random X-chromosome inactivation (XCI) occurring in female mammals. However, the *O* gene in the cat genome has not been identified, and the genetic variation responsible for the orange coloration remains unknown. We report here that a 5.1-kilobase (kb) deletion within an intron of the X-linked *ARHGAP36* gene, encoding a Rho GTPase activating protein, is closely and exclusively associated with orange coloration. The deleted region contains a highly conserved putative regulatory element, whose removal presumably cause altered *ARHGAP36* expression. Notably, *ARHGAP36* expression in cat skin tissues is linked to the suppression of many melanogenesis genes, potentially shifting pigment synthesis from eumelanin to pheomelanin. Furthermore, we find evidence that the gene undergoes XCI in female human and mouse cells, and XCI-dependent CpG island methylation consistent with random XCI in female domestic cats. The 5.1-kb deletion seems widespread in domestic cats with orange coat coloration, suggesting a single origin of this coat color phenotype.

## Introduction

The domestic cat (*Felis silvestris catus*) is a valued companion animal, a vermin-control agent, and a provider of important biomedical models, which most likely descended from a wildcat progenitor subspecies, *Felis silvestris lybica*, around 10,000 years ago^1,2,3^. Domestic cats display a diversity of phenotypic variation, including a range of coat coloration patterns resulting from interactions between multiple genetic loci, making them excellent models for studying gene function and regulation.

The X-linked *orange* (*O*) locus is one of the coat coloration loci, which controls an unknown molecular mechanism that causes the suppression of black-brownish pigmentation (eumelanin) and promotion of orange coloration (pheomelanin)^4^. Phenotypic variants of this locus can be seen in the tortoiseshell (mottled orange and black-brownish pattern) and the calico cats (mosaic pattern of large orange, black-brownish, and white patches), which are almost exclusively female. These coloration patterns arise as a result of X-chromosome inactivation (XCI), which is an important epigenetic mechanism that equalizes X-linked gene dosage between females and males. More than 60 years ago, it was suggested that one of the two X chromosomes in a female embryo is randomly selected and inactivated in each cell during early development, and that descendant cells maintain the same inactivation pattern as the ancestral cell^5^. As a result, the alternative expression of the *orange* allele versus the wildtype allele in different skin patches creates a mosaic coloration pattern in female cats heterozygous at the *O* locus^5^. Rare occurrence of the mosaic phenotype in males can be explained by sex chromosome aneuploidy (XXY), chimerism, mosaicism, or somatic mutations^6^.

While the coat color phenotypes of tortoiseshell and calico cats are often quoted as visually recognizable examples of XCI in female mammals, the gene and the genetic variation responsible for the orange coloration remain unknown. A previous genetic mapping study using microsatellite markers located the *O* locus within a 9.7-megabase (Mb) region of the cat X chromosome^7^. A more recent study using a 63K DNA array identified a 1.5-Mb haplotype block associated with orange coloration within this 9.7-Mb region^8^. The study also provided a list of twelve genes located within the haplotype region as the candidates for the *O* gene; however, none were known to regulate melanin synthesis.

In this study, we searched for the gene and genetic variation associated with the orange coloration by comparing the genomic sequences of cats with orange coloration, including tortoiseshell and calico cats, and control cats. We identified a 5.1-kilobase (kb) deletion in one of the protein-coding genes located within the haplotype region, which was closely and exclusively associated with orange coloration. We further present evidence that this gene undergoes XCI, most likely random XCI, in multiple mammalian species.

## Results

### Systematic screening for genetic variations responsible for orange coloration using whole-genome sequencing (WGS)

To identify the genetic variation responsible for the orange coloration and X-linked *O* gene in domestic cats, we sequenced the DNA samples from 18 cats, including eight calico cats, one tortoiseshell cat, one cat with both orange and white coat regions, and eight cats without orange coloration (Supplementary Tables 1 and 2, Extended Data Fig. 1A). We mapped the WGS reads to the AnAms1.0 cat genome, a high-quality chromosome-scale sequence assembly with only one short unmapped scaffold^9^. Thirteen putative protein-coding genes, overlapping with the twelve genes previously reported^8^, were identified in the 1.5-Mb haplotype region (Supplementary Table 3).

The AnAms1.0 genome was derived from a silver tabby American Shorthair^9^ (Extended Data Fig. 1A), and we used this sequence as a reference (wildtype) to identify genetic variations associated with the orange coloration. Within the 1.5-Mb haplotype region, we identified 2,569 single nucleotide polymorphisms (SNPs) in ten cats with the orange coloration, and 2,073 SNPs in eight cats without orange coloration. Removal of SNPs shared by the two groups identified 594 SNPs unique to the orange coloration group (Extended Data Fig. 2A). Using published WGS data from nine cats with color photographs, contributed by the 99 Lives Cat Genome Sequencing Initiative^10^ (Supplementary Table 4) (photographs available in the paper or at https://cvm.missouri.edu/research/feline-genetics-and-comparative-medicine-laboratory/99-lives/successfully-sequenced-cats/), we identified 901 SNPs in seven non-orange cats. After removing the SNPs shared with this pool, a total of 450 SNPs unique to the orange group remained, none of which were frameshift, missense, or nonsense mutations in the protein-coding exons (Extended Data Fig. 2B).

To search for other types of variations, we analyzed the sequence depth across the 1.5-Mb *O* haplotype region and identified a 5.1-kb deletion present only in the orange cat group (Fig. 1A). This deletion was located within the first intron of *ARHGAP36* and was surprisingly present in all ten sequenced cats with the orange coloration, but in none of the eight cats without orange coloration (Fig. 1A, Extended Data Fig. 3A). Notably, the eight calico cats and one tortoiseshell cat were heterozygous for the deletion, consistent with the mosaic coat color phenotype. Furthermore, analysis of the published data from the above-described nine cats with coat color information confirmed the presence of this deletion only in those with the orange coloration, namely a calico cat named Cali and her father, Orion^10^ (Extended Data Fig. 3B, Supplementary Tables 4 and 5). Again, the calico cat was heterozygous for the deletion. While we could not exclude at this stage that one of the identified SNPs was responsible for the *O* allele, the consistent and exclusive presence of the deletion made it a strong candidate for the causative genetic variation. Interestingly, the SNP (position 108,984,866 in AnAms1.0, or 110,230,748 in felCat9) (hereafter we use AnAms1.0 coordinate) previously found to be most closely associated with the orange coloration^11^ was located in an intron of *ENOX2*, a protein-coding gene located closest to *ARHGAP36* (Supplementary Table 3).

**Figure 1.**
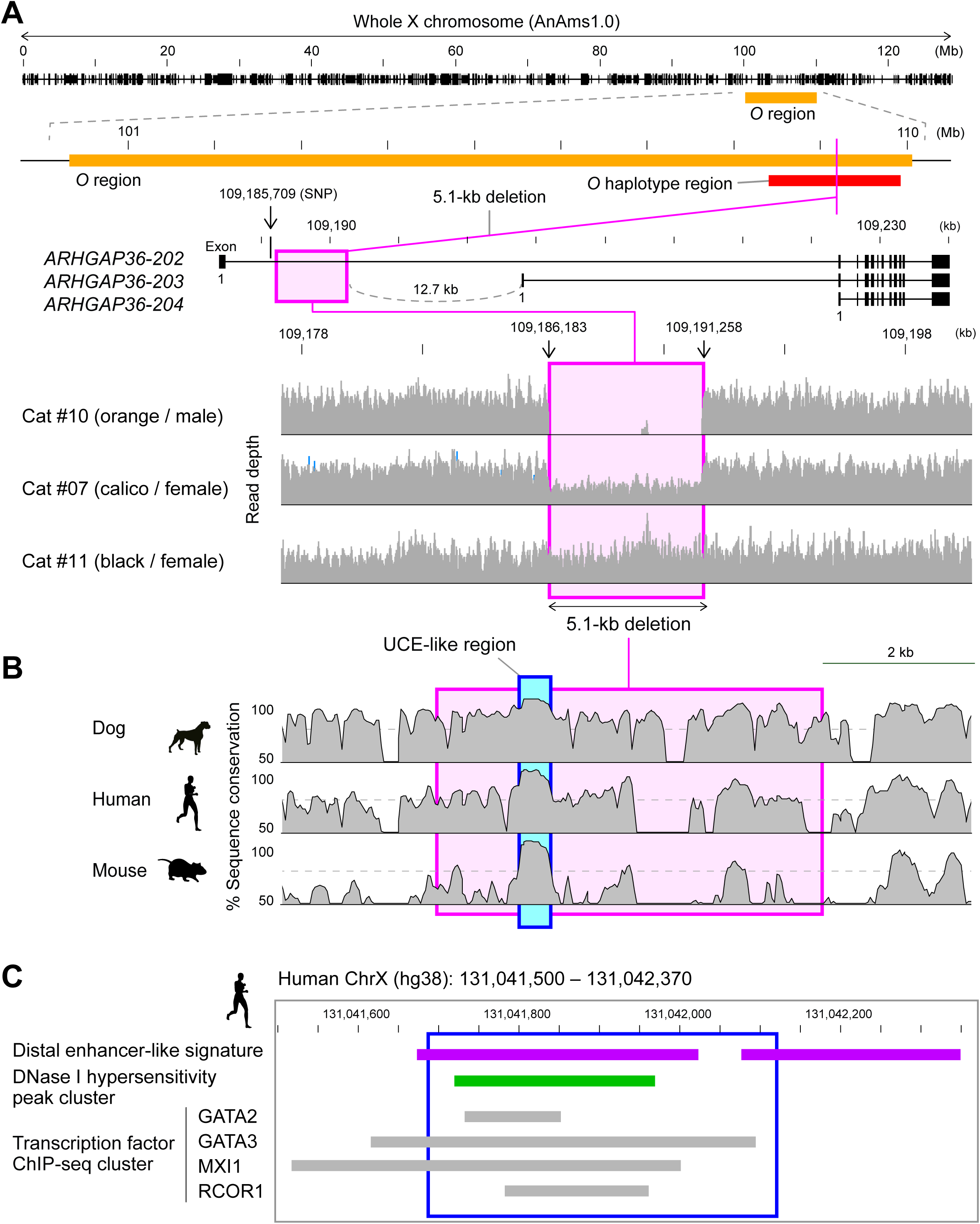
Identification of the 5.1-kb deletion within the *O* haplotype region. (A) Location of the 5.1-kb deletion associated with the orange coloration in the cat X chromosome. An orange bar indicates the 9.7-Mb *O* region identified using microsatellite markers^7^ and a red bar marks the 1.5-Mb *O* haplotype region identified using SNP microarray^8^. A pink box marks the 5.1-kb region deleted in cats with the orange coloration. Exon-intron structure of *ARHGAP36* is shown with exon usage generating three isoforms. Representative WGS read-depth profiles across the 5.1-kb region are shown at the bottom for three cats with indicated coat color and sex. (B) Sequence similarity profiles of the region containing the 5.1-kb deletion. Sequence similarity between domestic cats and indicated species is shown. A pink box highlights the deleted region, while a blue box marks the UCE-like sequence. (C) ENCODE annotations for the UCE-like sequence on human chromosome X. Shown are features of the hg38 131,041,688-131,042,122 region corresponding to the cat UCE-like sequence (boxed by blue lines) and adjacent regions, retrieved from the UCSC Genome Browser. Purple bars indicate regions showing distal enhancer-like signatures and a green bar highlights the DNase I hypersensitive site cluster identified in neuroblastoma. Gray bars represent ChIP-seq peak clusters for indicated transcription factors in neuroblastoma.

### Strong association between the deletion and orange coloration

To confirm the association between the 5.1-kb deletion within an intron of *ARHGAP36* and the orange coloration, additional DNA samples (n = 40) were analyzed by polymerase chain reaction (PCR). The study showed that all six calico cats, including one rare male calico cat (#24) (Extended Data Fig. 1B), and all five tortoiseshell cats had the deletion in one of the two X-chromosomes (Extended Data Fig. 3C, Supplementary Table 1). The fact that the male calico was heterozygous for the deletion is consistent with the presence of sex chromosome aneuploidy (presumably XXY)^6^. Three additional cats with the orange coloration but no black-brownish coloration (#30, #31, and #32) had this deletion in a hemizygous (male) or homozygous manner (female), while none of the non-orange cats did (n = 26) (Supplementary Table 1). Combining our WGS and PCR data, a total of 58 cats (24 orange and 34 non-orange) demonstrated a 100% link between the deletion and the orange coloration (Fisher’s exact test: *P* = 2.2e-16) (Table, Supplementary Table 1). Addition of the information from the publicly available WGS data of the above-described nine photographed cats further supported this link in a total of 67 cats (26 orange and 41 non-orange) (*P* = 2.2e-16).

Next, we sought to determine whether any of the 450 SNPs found only in the orange cat group were linked to the 5.1-kb deletion because such a SNP could also contribute to the phenotype. We found that only one SNP (C->T, at position 109,185,709), located 0.5-kb upstream of the deletion within an intron of *ARHGAP36* (Fig. 1A), was present in all ten sequenced cats of our orange group (Extended Data Fig. 2C,D). We then used 258 WGS datasets, mostly from the 99 Lives Cat Genome Sequencing Initiative^10^ (including those from the nine cats with color images), and found that, while all 61 cats carrying the 5.1-kb deletion (21 hemizygous males, 12 homozygous females, and 28 heterozygous females) had the SNP, none of the 197 cats without it did (Supplementary Tables 4 and 5). All 28 females heterozygous for the deletion were also heterozygous for the SNP. These results indicate that the deletion and a linked SNP are present within an intron of *ARHGAP36* in cats with the orange coat coloration.

### Ultraconserved element (UCE)-like sequence in the deleted region

We then performed a more detailed analysis of the deletion and SNP. Sanger sequencing of the PCR products confirmed a 5,076-nucleotide (nt) deletion spanning from position 109,186,183 to 109,191,258 (Extended Data Fig. 3D). The terminal ‘AA’ dinucleotide likely served as an acceptor template. Importantly, the deleted region contained a 436-nt sequence showing strong evolutionary conservation reminiscent of UCE (ref. 12) (Fig. 1B, Extended Data Fig. 3E). According to the data from the human ENCODE project^13^, this UCE-like sequence contained a DNase I hypersensitive site cluster in human neuroblastoma cells and chromatin immunoprecipitation-sequencing (ChIP-seq) clusters for transcription factors including GATA2/3, MXI1, and RCOR1 (Fig. 1C). Interestingly, while GATA family proteins are transcriptional activators^14^, MXI1 (Max interactor 1, dimerization protein) and RCOR1 (REST corepressor 1 or CoREST) are transcriptional repressors^15,16^. Therefore, this putative *cis*-regulatory element could function as a transcriptional enhancer or a repressor. In contrast, the SNP located about 0.5-kb upstream of the deletion (position 109,185,709) was not in a region conserved among species and therefore unlikely to have a role in gene regulation. It is likely that the 5.1-kb deletion removed a critical regulatory region, leading to altered expression of a nearby gene(s) such as *ARHGAP36*.

### Negative correlation between *ARHGAP36* and melanogenesis gene expression

We hypothesized that the *O* gene would show differential expression depending on the presence/absence of the deletion in skin tissues, especially in orange and black-brownish coat regions. We first performed RNA sequencing (RNA-seq) on skin tissues obtained from the auricles of different adult calico cats (#19, #20, and #22, females, precise ages unknown) (Supplementary Table 2). Of the thirteen protein-coding genes found in the *O* haplotype region, only *ARHGAP36* showed differential expression in different coat color regions, with its expression detected only in the orange region (Extended Data Fig. 4A). We then analyzed multiple skin tissues obtained from a young calico cat that died of unknown causes (#01, female, three weeks old) (Extended Data Fig. 1C, Supplementary Table 2). Of the genes in the *O* haplotype region, *ARHGAP36* and *IGSF1* showed differential expression (Fig. 2A). *ARHGAP36* was persistently expressed in the orange regions but variably expressed in the black-brownish regions (Fig. 2A, Extended Data Fig. 4A). One of the two black-brownish regions showed virtually no *ARHGAP36* expression. *IGSF1*, a gene located next to *ARHGAP36*, behaved similarly, suggesting that the two genes might share a regulatory mechanism, although the expression level of *IGSF1* was extremely low (< 0.6 fragments per kilobase of exon per million reads mapped [FPKM]) (Fig. 2A). Notably, *ARHGAP36-203* was the only mRNA isoform produced in the skin samples (Extended Data Fig. 4B). Since the expression pattern observed in the black-brownish regions should be the wildtype pattern, the more persistent *ARHGAP36*, and possibly *IGSF1*, expression induced by the deletion could shift the pigment synthesis from eumelanin to pheomelanin in the orange regions.

**Figure 2.**
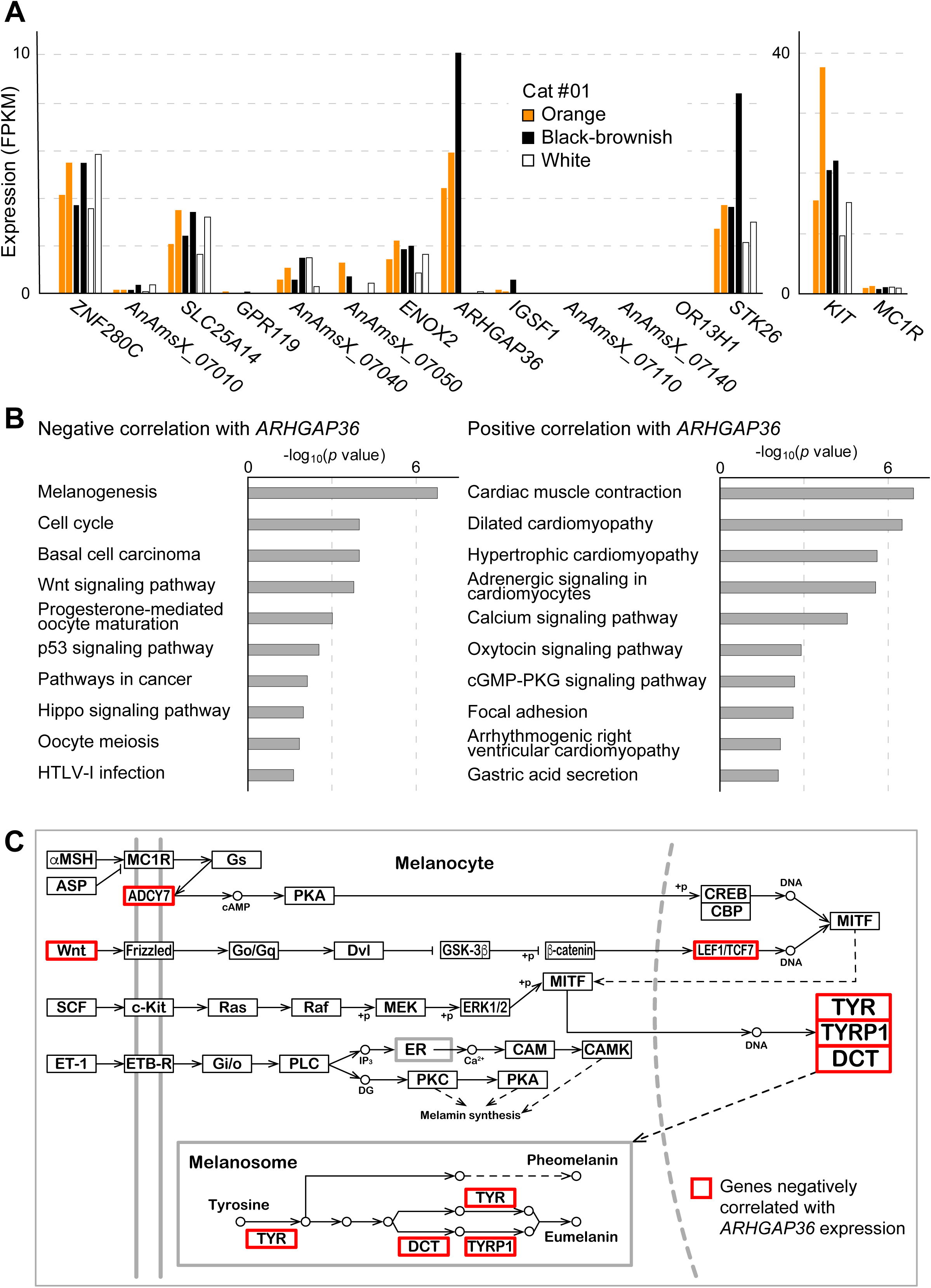
Negative correlation between *ARHGAP36* and melanogenesis gene expression. (A) Expression levels of the thirteen protein-coding genes of the *O* haplotype region in cat skin tissues. RNA-seq data were obtained in skin regions of different coat colors from a calico cat (#01). Expression levels of key melanocyte genes, *KIT* and *MC1R*, are also shown. (B) Top ten KEGG pathways in which genes negatively and positively correlated with *ARHGAP36* are enriched. The pathways are ranked by significance. (C) KEGG diagram of the melanogenesis pathway highlighting genes negatively correlated with *ARHGAP36* expression (red boxes). The original KEGG pathway diagram was slightly modified for clarity.

Based on the publicly available gene expression databases, such as the Genotype-Tissue Expression (GTEx)^17^, both *ARHGAP36* and *IGSF1* are highly expressed in neuro-endocrinological tissues, such as the hypothalamus, pituitary gland, and adrenal gland. Notably, these tissues are rich in neural crest-derived cells, from which melanocytes also originate^18^. *IGSF1* encodes a member of the immunoglobulin superfamily, and loss-of-function mutations in this gene cause X-linked central hypothyroidism and testicular enlargement (OMIM300888)^19^. There is currently no known role for *IGSF1* in hair follicle development or melanogenesis. In addition, its expression was very low in cat skin tissues as described above (Fig. 2A, Extended Data Fig. 4A). *ARHGAP36* encodes a member of the Rho GTPase-activating protein family, but a systematic study has shown that this protein lacks the GTPase-activating activity^20^. Rather, it has been reported that ARHGAP36 mediates and activates Hedgehog (HH) signaling and suppresses protein kinase A (PKA) signaling^21–25^. Importantly, the HH and PKA signaling pathways mediate hair follicle development and melanogenesis, respectively, and dysregulation of human *ARHGAP36* is implicated in X-linked Bazex-Dupré-Christol syndrome (BDCS, OMIM301845)^26^, characterized by follicular atrophoderma, congenital hypotrichosis, and multiple basal cell naevi and carcinomas. Hence, its known biological roles make *ARHGAP36* an excellent candidate for the *O* gene.

We then sought genes that were positively and negatively correlated with *ARHGAP36* expression in the skin samples from orange and black-brownish regions, excluding white regions where no melanogenesis is expected. We found 341 genes showing positive correlation with *ARHGAP36* and 517 genes showing negative correlation (Supplementary Table 6). Strikingly, an analysis using the Kyoto Encyclopedia of Genes and Genomes (KEGG) in the David system^27,28^ revealed that the negatively correlated genes were enriched in the melanogenesis pathway (Fig. 2B). Examples of the melanogenesis genes negatively correlated with *ARHGAP36* included *ADCY7*, *DCT*, *LEF1*, *TCF7*, *TYR*, *TYRP1*, *WNT10B*, *WNT3A*, *WNT5A*, and *WNT6*. The roles of the protein products were distributed in the whole melanogenesis pathway, from ligands activating the signaling pathways to enzymes directly involved in the melanin synthesis (Fig. 2C). Notably, among the melanin synthesis enzymes, TYRP1 and DCT are involved in eumelanin synthesis but not pheomelanin synthesis. Therefore, their downregulation may attenuate eumelanin synthesis and cause a shift towards pheomelanin synthesis^29^. The negative correlation between *ARHGAP36* and melanin synthesis enzyme genes was also seen in the auricle skin samples from different calico cats (Extended Data Fig. 4C).

### *ARHGAP36* is subject to XCI in female mammalian cells

The mosaic coat color of tortoiseshell and calico cats is attributed to the random inactivation of the X-linked *O* gene in *O*/*o* female cells^5^. However, a proportion of X-linked genes is known to escape XCI to varying degrees^30^. We therefore wished to examine whether *ARHGAP36* of the domestic cat is subject to random XCI in female cells and whether melanocytes in black-brownish and orange regions express the different alleles. To do this, we would need tissue samples from female cats (preferably from tortoiseshell or calico cats) carrying informative SNPs within the *ARHGAP36* transcripts. However, we have not yet had a chance to access such samples.

The XCI is investigated in detail in humans and mice. One of the authors (T.S.) previously investigated the inactivation of the X-linked genes using fibroblast cells derived from female mouse embryos obtained by crossing a JF1 female and a C57Bl/6J male^31,32^. In the study, the X chromosome from C57Bl/6J was fixed to be active by an internal deletion of *Xist* (X^B6-ΔA^), a gene essential for XCI^31^. We reprocessed the RNA-seq data and found that all 172 SNP-containing *Arhgap36* reads were from the X^B6-ΔA^, indicating that the gene is subject to XCI in mice. We also reprocessed the chromatin RNA-seq data from a similar study using embryonic fibroblast cells where the X chromosome from *Mus musculus castaneus* was genetically fixed to be inactive^33^ and found that all *Arhgap36* transcripts were from the other X (27 reads containing informative SNPs). Regarding the XCI of human *ARHGAP36*, a SNP-based allelic imbalance analysis was performed for a total of 409 X-linked genes in multiple lymphoblastoid and fibroblast cells^34^. The data indicated that human *ARHGAP36* (designated *FLJ30058* in that study) was subject to XCI in all ten female cell-lines possessing informative SNPs. These results show that, while direct evidence is lacking for domestic cats, *ARHGAP36* is subject to XCI in multiple mammalian species.

### DNA methylation status of cat *ARHGAP3*6 is consistent with random XCI

DNA methylation can serve as a hallmark for genes that are subject to XCI, as X-linked CpG islands (CGIs), especially promoter-associated ones, are highly methylated on the inactive X but almost unmethylated on the active X (ref. 35). Using published whole-genome bisulfite sequencing (WGBS) datasets from human and mouse tissues^36,37^, we first compared the methylation levels of the *ARHGAP36* CGIs between the sexes, which showed significantly higher methylation levels in females of both species (Extended Data Fig. 5A,B), consistent with their XCI-dependent methylation. We then investigated the published allelic methylation data from hybrid mouse embryonic fibroblast cells carrying a *Mus musculus castaneus*-derived inactive X (ref. 33) and found that the *Arhgap36* CGI is more methylated on the inactive X (Extended Data Fig. 5A).

To determine whether the CGIs of cat *ARHGAP36* undergo XCI-dependent methylation, we performed WGBS on skin tissues of different colors from a female calico cat (#01) and peripheral blood cells from two other cats (female #04 and male #10) (Supplementary Table 2). We also obtained publicly available WGBS data from a male cat named Boris^38^. Globally, X-linked promoter CGIs (n = 362) were more methylated in females, ranging from 20% to 40%, than in males, ranging from 0% to 5% (Fig. 3A), consistent with the XCI-dependent methylation. We then looked at the *ARHGAP36* CGIs and found higher methylation in females than in males, just as the most other X-linked CGIs (Fig. 3B). In contrast, the CGIs of known XCI-escaper genes, such as *KDM6A, MED14,* and *DDX3X* (ref. 39), were unmethylated in both sexes (Extended Data Fig. 5C). We then analyzed the allelic methylation status of one of the *ARHGAP36* CGIs where two SNPs were available (A/G and C/A substitutions at positions 109,204,156 and 109,204,409, respectively). The result showed that both alleles are moderately methylated (Fig. 3C), consistent with the random XCI in tissues consisting of a heterogeneous cell population, rather than parental-origin-dependent (imprinted) or other types of skewed XCI (ref. 40). Altogether, our results corroborate Lyon’s hypothesis that the color patterns of tortoiseshell and calico cats are the results of random XCI of the *O* gene^5^.

**Figure 3.**
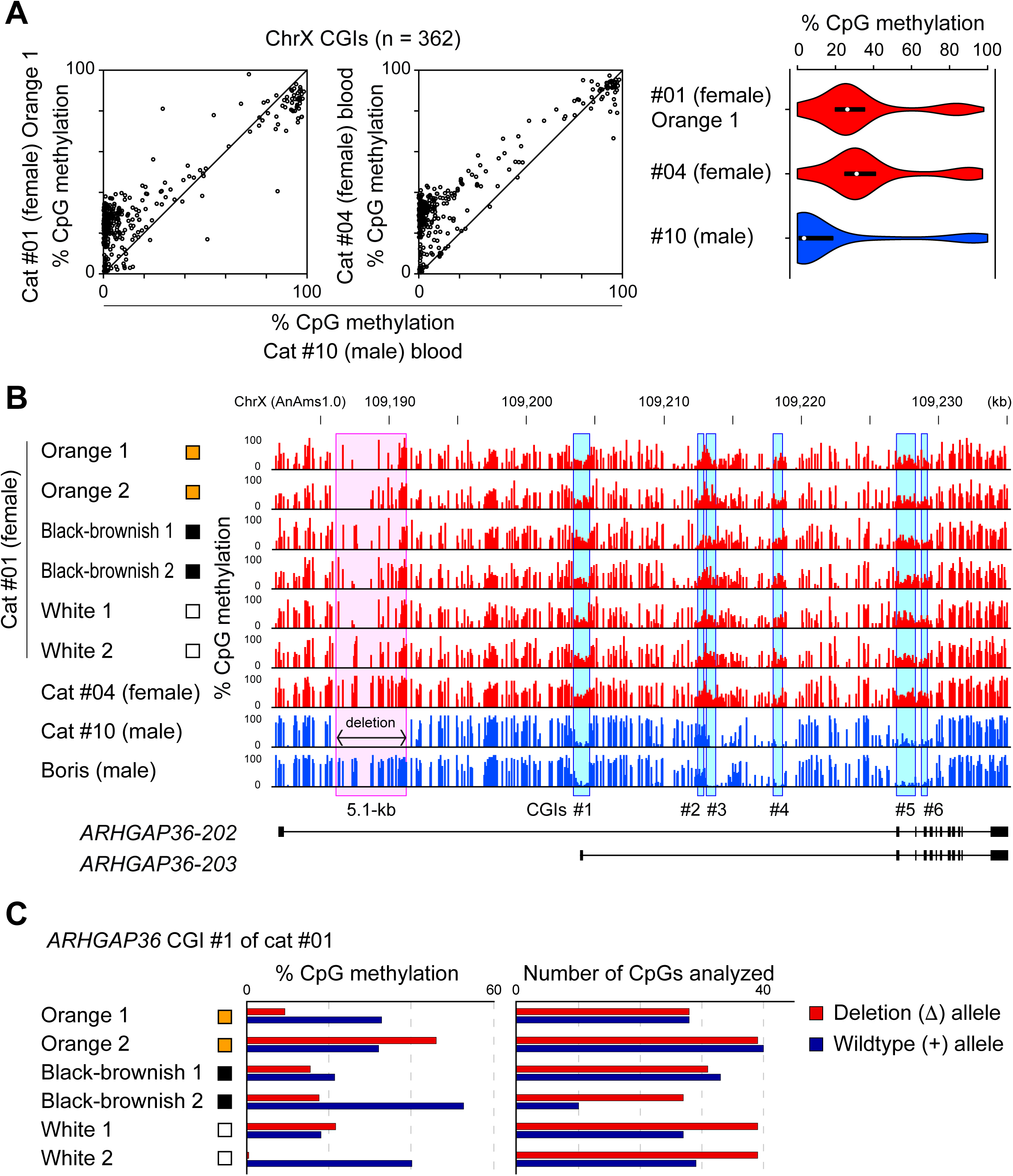
DNA methylation profiles of the X chromosome and *ARHGAP36* in female and male cats. (A) Scatter plots and violin plots comparing CpG methylation levels of 362 X-linked promoter CGIs between female and male cats. The WGBS data were from skin (cat #01, orange patch 1) and peripheral blood cells (cats #04 and #10). (B) CpG methylation profiles of the cat *ARHGAP36* region in female (red) and male cats (blue). The WGBS data were obtained from skin regions (cat #01) and peripheral blood cells (cats #04 and #10). The WGBS data of peripheral blood cells from a male cat named Boris was obtained from a publicly available dataset (SRX548709). A pink box indicates the region corresponding to the 5.1-kb deletion, and blue boxes indicate the CGIs. (C) Allelic CpG methylation of CGI #1 of *ARHGAP36*. The CpG methylation levels (left) and the number of CpGs analyzed using SNP-containing WGBS reads (right) are shown. The data were from different skin regions of a calico cat (#01). Two SNPs were used to analyze allele-specific reads: *cis* association of the SNPs with the deletion (Δ) was determined using the data from cats hemizygous (males) and homozygous (females) for the deletion. Note that CGI #1 overlaps with the most active promoter in the skin (Extended Data Fig. 4B).

### Origins of the genetic variations responsible for the coat colors

As described above, all 24 cats with orange coloration that we analyzed had the 5.1-kb deletion (Table, Supplementary Table 1). They were either owned or feral cats, which were mostly from the Fukuoka City area of Japan. Also, sixty-one of the 258 cats contributed by the 99 Lives Cat Genome Sequencing Initiative^10^ (Supplementary Table 4), which had exactly the same deletion, included those from the United States, Europe, and the Middle East^41^. Thus, it is likely that the 5.1-kb deletion is widely distributed in domestic cats worldwide, suggesting the single origin of the orange coat color phenotype.

Calico cats have white coat regions in addition to orange and black-brownish patches. A feline endogenous retrovirus insertion within the *KIT* gene on chromosome B1 has been identified as the causative genetic variation for white spotting^42^. To detect the white spotting allele (*w^s^*) in our cat samples, we performed a PCR assay. All cats with white coat regions (n = 25), including all fourteen of the calico cats, carried at least one *w^s^* allele (Supplementary Table 1), consistent with a previous report^42^. In contrast, cats without white coat regions (n = 33), including all six tortoiseshell cats, did not carry this allele (Supplementary Table 1). Altogether, the genetic basis of the coat color patterns of calico and tortoiseshell cats is likely to be common worldwide.

Lastly, the golden hamster (*Mesocricetus auratus*) exhibits orange coloration due to the activity of an X-linked gene designated *Sex-linked yellow* (*Sly*)^43^. Using a high-quality chromosome-scale sequence assembly of the golden hamster genome^44^, we examined whether the genomic position of *Arhgap36* corresponds to the genetic location of *Sly*. This analysis showed that the two loci are located in different regions of the X chromosome (Extended Data Fig. 6), supporting the previous conclusion reached by others that *Sly* and *O* are distinct^43^. Together with the knowledge obtained so far in humans and mice, this result suggests that the genetic contribution of *ARHGAP36* and the UCE-like sequence to orange coloration may be unique to certain mammalian species.

## Discussion

More than 60 years ago, Lyon suggested that the alternative black-brownish and orange patches of tortoiseshell cats are consistent with the random inactivation of the heretofore unidentified X-linked *O* gene in heterozygous females^5^. Our systematic screening now identifies a 5.1-kb deletion within an intron of *ARHGAP36*, which is among the previously listed candidate genes in the *O* haplotype region^8^, as the genetic variation responsible for the orange coloration. Although no non-invasive method is available at present for direct demonstration of the causal relationship in this companion animal species, a perfect association between the existence of the identified genetic variation deleting a putative regulatory element and the orange coat coloration strongly supports our conclusion. While *ARHGAP36* and its adjacent gene, *IGSF1*, appear to share some regulatory mechanisms, the observed higher expression of the former in the skin, its known roles in signaling pathways related to hair follicle development and melanogenesis (the HH and PKA pathways), and its implications in human skin diseases involving hair follicles and naevi^26^ all point to it as an excellent candidate for the long-sought *O* gene. Furthermore, we provide evidence that *ARHGAP36* is subject to XCI in female human and mouse cells. Our results on the allelic DNA methylation status of the *ARHGAP36* CGIs are also consistent with the random inactivation of this gene in female cats.

*ARHGAP36* encodes a member of the Rho GTPase activating protein family and is implicated in neuronal development, bone formation, hair follicle development, and cancer development or progression through positive and negative regulation of the HH and PKA signaling pathways, respectively^21–26^, but no role has been reported in melanogenesis. In fact, no locus affecting hair or coat color has been mapped at the orthologous location in humans or mice. Additionally, the location of the X-linked *Sly,* which is responsible for the orange coat coloration of the golden hamster, is distinct from both the *O* locus^43^ and the *ARHGAP36* gene (this study). Thus, ARHGAP36 protein is a novel factor regulating the melanogenesis pathway, and this role may be specific to the domestic cat or the feline family. Consistent with this, our attempts to generate mouse models by introducing the genetic variation have not been successful: neither a deletion corresponding to the one found in domestic cats nor a more targeted deletion restricted to the UCE-like sequence caused no discernible coat color change in several engineered mouse lines.

How does *ARHGAP36* regulate melanogenesis? According to the public databases, *ARHGAP36* is highly expressed in tissues containing neural crest derivatives, but it is unknown whether the melanocyte lineage, which is also a neural crest derivative, are the primary site of *ARHGAP36* expression in the skin. A previous immunostaining study however reported the detection of *ARHGAP36* expression in a small number of hair follicle cells in the outer root sheath of healthy human skin^26^. Since our study using commercially available antibodies did not work for cat skin samples, the precise site of *ARHGAP36* expression remains an open question. In any case, our RNA-seq data shows that *ARHGAP36* expression in melanocytes, or cells communicating with them, negatively regulates a variety of melanogenesis pathway genes including those encoding signaling molecules (ligands) and enzymes directly involved in melanin synthesis. It is interesting that ARHGAP36 protein appears to lack the GTPase activating activity^20^ but has the potential to inhibit PKA signaling directly by blocking the catalytic activity of PKA or by accelerating ubiquitin-mediated degradation of the catalytic subunit of PKA^22^. The PKA signaling elicited by MC1R bound to melanocyte stimulating hormone is a fundamental pathway regulating melanin synthesis^45^ and, in fact, genetic variations in MC1R are associated with red hair and fair skin of humans (increased pheomelanin and decreased eumelanin)^46^. So, if *ARHGAP36* is expressed in melanocytes, it could directly modulate the PKA pathway to alter the balance between eumelanin and pheomelanin.

The genomic region of the 5.1-kb deletion contains a UCE-like sequence containing putative binding sites for transcriptional activators and repressors. Based on the expression patterns of cat *ARHGAP36* in colored coat regions, we speculate that the deletion disrupts its regulated expression and causes inappropriate activation. However, we do not know the precise change caused by the deletion since the follicle melanogenesis is a complex process regulated by a variety of intrinsic and extrinsic factors. For example, it is restricted to the anagen stage of the hair cycle and coupled to the life cycle of melanocytes with changes in their distribution and differentiation^45^. Therefore, we would first need to know the precise site and timing of *ARHGAP36* expression in healthy cat follicles. Since samples required for such a systematic study are not easily accessible in this non-experimental animal species, we are hoping that recently derived feline induced pluripotent stem (iPS) cells^47^ and their use in constructing in vitro systems including organoids would resolve some of the problems. In particular, derivation of naïve female iPS cells carrying two active X chromosomes and their differentiation into melanocytes and other cells would recapitulate the random XCI and production of different pigment species in different cells. Further investigation warrants detailed understanding of how the deletion eventually regulates pheomelanin synthesis.

In conclusion, we have identified the X-linked *ARHGAP36* gene as the strong candidate as the long-sought *O* gene of the domestic cat. Although it is still unclear how the identified deletion switches eumelanin to pheomelanin, the variation likely dominates the cat population with orange coloration. We also provide evidence that *ARHGAP36* is most likely subject to random XCI in the domestic cat as previously suggested by Lyon^5^. How this gene became employed in melanogenesis in certain species and how it exerts pleiotropic biological effects in mammals are interesting future questions.

**Table.** Association between orange coloration and a 5.1-kb deletion within *ARHGAP36*.

| Coat color pattern | Genotype regarding the deletion* |  |  |  |  |  |
| --- | --- | --- | --- | --- | --- | --- |
|  | Female |  |  | Male |  |  |
| | +/+ | $\Delta$/+ | $\Delta/\Delta$ | +/Y | $\Delta$/Y | $\Delta$/+/Y |
| Calico | 0 | 13 | 0 | - | - | 1 |
| Tortoiseshell | 0 | 6 | 0 | - | - | - |
| Orange | 0 | 0 | 2 | 0 | 2 | - |
| Non-orange | 16 | 0 | 0 | 18 | 0 | - |
| Total | 16 | 19 | 2 | 18 | 2 | 1 |

\* $\Delta$ , 5.1-kb deletion; +, wildtype; Y, Y chromosome.

## Methods

### Ethics statement

Sampling of domestic cats was carried out in accordance with the ethical guidelines of Kyushu University using the protocols approved by the Animal Experiment Committee (#A21-433-1). Blood samples and small tissue pieces were acquired by veterinarians during routine veterinary practice or spaying/neutering surgery. Skin tissues from an animal that died of unknown causes were voluntarily donated by a breeder.

### DNA and RNA extraction

Genomic DNA was extracted from whole blood cells using the NucleoSpin Blood kit (Macherey-Nagel) or from tissue samples by a standard method involving proteinase K digestion and isopropanol precipitation. Total RNA was extracted from tissue samples using TRIzol Reagent (Thermo Fisher Scientific).

### WGS and data analysis

Libraries were prepared from genomic DNA using the TruSeq DNA PCR-Free LT Library Prep Kit (Illumina) and sequenced to generate 53-nt paired-end reads on an Illumina NovaSeq 6000 equipped with NVCS v1.6.0 and RTA v3.4.4. The reference genome of the domestic cat (AnAms1.0) was obtained from the Cats-I database^9^. All paired-end reads were aligned to the reference genome using the Burrows-Wheeler Aligner (BWA v0.7.17)^48^. After removing duplicate reads, single nucleotide substitutions and small insertion-deletion variants (indels) were identified. Variant Call Format files were generated using the Genome Analysis Toolkit (GATK v4.1.3)^49^. These files were annotated using SnpEff v4.3 (ref. 50) to assess the effect of variants on gene function.

Raw fastq files from published WGS datasets of 258 cats, including nine with clear color images, contributed primarily by the 99 Lives Cat Genome Sequencing Initiative^10,41^, were obtained from the Sequence Read Archive. The datasets were analyzed using our proprietary pipeline. Human (hg38 and hg19) and mouse (mm10) X chromosome sequences and ENCODE data were downloaded from the UCSC Genome Browser. Sequence alignment between multiple mammalian species was performed using mVISTA (ref. 51) and ClustalW v2.1 (ref. 52). The golden hamster genome sequence was obtained from the Sequence Read Archive (DRA010879)^44^.

### Genotyping

The cats were genotyped for alleles of *ARHGAP36* (ϑ′/+) and *KIT* (*w*^+^/*w^s^*) by PCR using KOD One PCR Master Mix (TOYOBO) with primers listed in Supplementary Table 7. Some animals were also examined for the presence of *SRY* on the Y chromosome.

### Sanger sequencing

Sanger sequencing of PCR products was performed to determine the exact position and size of the deletion using BigDye Terminator v3.1 (Thermo Fisher Scientific) and primers listed in Supplementary Table 7.

### RNA-seq and data analysis

Libraries were prepared from total RNA using NEBNext rRNA Depletion Kit, NEBNext Ultra II Directional RNA Library Prep Kit for Illumina, and NEBNext Multiplex Oligos for Illumina (96 Unique Dual Index Primer Pairs) (NEB). Sequencing was performed using a SP Reagent Kit (Illumina) on an Illumina NovaSeq 6000 equipped with NVCS v1.6.0 and RTA v3.4.4 to generate 53-nt paired-end reads. Adapter sequences and low-quality bases were removed from the 5’ and 3’ ends, respectively. Reads were aligned to the AnAms1.0 genome by HISAT2 v2.0.5 (ref. 53). Transcripts were assembled using StringTie v2.1.4 (ref. 54). To characterize the gene regulatory network involving *ARHGAP36*, we identified genes that showed positive and negative correlation with *ARHGAP36* expression in skin samples from a calico cat (#01). Only the orange and black-brownish regions (n = 2 each) were used for the analysis because we did not expect melanogenesis in the white regions. (Indeed, we observed no *ARHGAP36* expression in the white region.) We then grouped the four datasets into two, *ARHGAP36* positive (two orange and one black-brownish samples) and *ARHGAP36* negative (one black-brownish sample). Then, a gene was considered to be positively correlated if its FPKM value was greater than 1 and at least five times higher in all three *ARHGAP36*-positive samples compared with the *ARHGAP36*-negative sample. Conversely, a gene was considered to be negatively correlated if its FPKM was greater than 1 and at least five times higher in the *ARHGAP36*-negtive sample compared to the *ARHGAP36*-positive samples. Then, genes positively and negatively correlated with *ARHGAP36* were subjected to the KEGG pathway analysis using DAVID^27,28^. RNA-seq data from mouse embryonic fibroblasts (GSE112097 and GSE121184)^31,33^ were reprocessed as described in the original manuscript for SNP-based allelic expression analysis of mouse *Arhgap36*. RNA-seq data from various cat organs and tissues were acquired from the publicly available dataset (PRJNA312519) and processed using the aforementioned pipeline.

### WGBS and data analysis

Genomic DNA spiked with 2 ng lambda phage DNA (Promega) was subjected to bisulfite conversion. WGBS libraries were prepared using the post-bisulfite adapter tagging method^55^. Sequencing was performed using a S1 Reagent Kit (Illumina) on an Illumina NovaSeq 6000 equipped with NVCS v1.6.0 and RTA v3.4.4 to generate 108-nt single-end reads. Adapter sequences and low-quality bases were removed from the 5’ and 3’ ends, respectively. Reads were aligned to the AnAms1.0 genome using Bismark v0.10.0 (ref. 56) with a seed length of 28, allowing a maximum of one mismatch in the seed and enabling the “--pbat” option. Only uniquely aligned reads were used in the subsequent analysis. Data from both DNA strands were combined. Bisulfite conversion rate was estimated using reads uniquely aligned to the lambda phage genome. Allele-specific CpG methylation levels were calculated by using SNPsplit v0.3.2 (ref. 57), based on a list of SNPs identified in *ARHGAP36*. Publicly available WGBS datasets were obtained from the Sequence Read Archive for a male cat named Boris (SRX548709)^38^, human cytotrophoblasts (NBDC No. hum0086)^37^, and mouse liver (GSE106379)^36^. These data were processed using the same pipeline as the cat data analysis. In addition, pre-calculated bedGraph files on DNA methylation of mouse embryonic fibroblasts were obtained from the NCBI Gene Expression Omnibus (GSE122094)^33^. CGIs were predicted in the AnAms1.0 genome sequence using the method described by Gardiner-Garden and Frommer^58^.

## Data availability

All sequencing data related to this study are publicly available in the DDBJ/ENA/NCBI databases under BioProject No. PRJDB18200.

## Acknowledgements

We thank Chaoqing Wen, Junko Oishi, Hiroaki Ohishi, Sangwan Kim, Sayaka Misumi, Masanobu Deshimaru, Yoshihiro Izumi, and Takeshi Bamba for technical assistance, Sei Matsumura and Keiko Akenaga for their help in collecting the cat samples, and Shingo Hatoya, Hiroaki Yamamoto and Tomohisa Hirobe for valuable discussion. We also thank Naohito Nozaki, Yasuaki Mohri and Emi Nishimura for their technical advice, and the Common Research Facility/Research Promotion Unit of the Medical Institute of Bioregulation, Kyushu University, for technical support. This research project was partly supported by 619 crowdfunding backers on the READYFOR platform (https://readyfor.jp/projects/calico60).

## Author contributions

M.U., M.M., and H.S. conceived the study and planned the experiments. M.U., Y. Matsumoto, Y. Matsumura, M.M. collected the cat samples. W.K.A.U., M.U., Y. Miki, and M.M. performed the experiments and H.T., W.K.A.U., Y.N., and H.S. analyzed the data. Y.B. and T.S. analyzed the data from mouse studies. Y.B., Y.N., M.M., and H.S. supervised the entire project. H.T., W.K.A.U., M.U., M.M., and H.S. interpreted the results and wrote the paper with input from all authors.

## Additional information

Extended data is available for this paper.

Supplementary information The online version contains supplementary material.

