## Extended Data Figures for "A deletion at the X-linked *ARHGAP36* gene locus is associated with the orange coloration of tortoiseshell and calico cats"

**The PDF file includes:**
Extended Data Figures 1 to 6

**Other Supplementary Materials for this manuscript include the following:**
Supplementary Tables 1 to 7

**Extended Data Figure 1. Color images of donor cats.**

(**A**) Selected color images of cats analyzed using WGS. An image on the left is of Senzu, a female American Shorthair, whose genome sequence served as the reference in this study.

(**B**) Image of Miketa, a rare male calico cat (17 years old) analyzed using PCR.

(**C**) Source of the skin tissues used for RNA-seq analysis. Sampling was done from indicated color regions of a calico cat of 3 weeks old (cat #01) that died in a breeder's facility.

**Extended Data Figure 2. Analysis of SNPs in the *O* haplotype region.**

(**A**) Stepwise filtration of SNPs associated with the orange coloration. All SNPs identified in the *O* haplotype region of cats with the orange coloration (SNP group 1) were filtered twice to obtain more specific ones (SNP groups 3 and 5). Seven non-orange cats contributing to SNP group 4 (Monkey, Marcus, Aries, Dragon, Frank, Camilla, and Bobble) are identified using the color images in the 99 Lives Cat Genome Sequencing Initiative website.

(**B**) Number of SNPs mapped in each genomic feature. All SNPs identified in the *O* haplotype region (SNP group 1) and those that survived after the two filtration-steps (SNP groups 3 and 5) were analyzed. For the SNPs mapped in the coding regions, numbers are provided according to the mode of codon change.

(**C**) Number of the survived SNPs mapped in each feature of the indicated gene. Of the 450 SNPs of group 5, one hundred and forty were present in genic regions (a total of eight genes). A table at the bottom shows the number of the genic SNPs carried by each of the ten sequenced cats with the orange coloration (#01 to #10). Note that *ARHGAP36* is the only gene where a SNP(s) could be found in all orange cats.

(**D**) Presence/absence of *ARHGAP36* SNPs in each of the ten orange cats. The SNP at position 109,185,709 was the only one carried by all ten cats.

**Extended Data Figure 3. Characterization of the 5.1-kb region responsible for the orange coloration.**

(**A**) Comparison of the WGS read depth of the 5.1-kb region in the orange and non-orange cats. The ratio of the depth of the 5.1-kb region to that of the entire X chromosome is shown. F, female; M, male.

(**B**) Representative read depth profiles of publicly available WGS data from three cats with color images (a calico cat called Cali and her parents). A pink box marks the 5.1-kb region. The WGS data were from BioProject number PRJNA343385. The images are available in a published paper[10] or at the University of Missouri 99 Lives Cat Genome Sequencing Initiative website. F, female; M, male.

(**C**) Representative PCR results regarding the presence/absence of the deletion. Bands representing the wildtype and the deleted allele are marked by + and Δ, respectively. PCR was done with primers *ARHGAP36* F1 and R1, designed to flank the 5.1-kb region (Supplementary Table 7). The results shown at the bottom include sexing by detection of *SRY*. F, female; M, male.

(**D**) Sequence of the edges of the 5.1-kb deletion. Bold 'AA' nucleotides are the putative acceptor template for DNA recombination.

(**E**) Sequence alignment of the UCE-like sequence from different vertebrate species. Nucleotides matching the cat reference sequence are cased in black (UCE-like sequence) or blue (flanking regions).

**Extended Data Figure 4. Expression of the *O* region genes, *ARHGAP36* isoforms, and melanogenesis genes in different cat samples.**

(**A**) Expression levels of the thirteen protein-coding genes located in the *O* haplotype region in auricle from different cats. RNA-seq was performed on small pieces of auricles of different coat colors obtained from indicated cats. Expression levels of key melanocyte genes, *KIT* and *MC1R*, are also shown.

(**B**) Expression pattern of *ARHGAP36* isoforms in indicated tissues and organs, highlighting differences in exon usage. The RNA-seq data of our own (skin) and from publicly available datasets (auricle, spinal cord, embryo body, and hippocampus) (PRJNA312519) are used for the analysis. Schematic of the isoform structure is shown at the bottom, with the position of the 5.1-kb deletion (pink box).

(**C**) Expression levels of melanogenesis genes identified by the KEGG analysis in auricles of different coat colors from indicated cats. Expression levels of key melanocyte genes, *KIT* and *MC1R*, are also shown.

**Extended Data Figure 5. DNA methylation profiles of *ARHGAP36* in mice and humans and other X-linked genes in domestic cats.**

(**A**) CpG methylation profiles of the *Arhgap36* region in female (red) and male mice (blue). The CGI is highlighted by a blue box. The WGBS data were from mouse liver (GSE106379) and embryonic fibroblasts (MEF) (GSE122094). The latter was used for allelic methylation analysis on the active and inactive X.

(**B**) CpG methylation profiles of the human *ARHGAP36* region in females (red) and males (blue). The CGIs are highlighted by blue boxes. The WGBS data were from human cytotrophoblasts (NBDC No. hum0086).

(**C**) CpG methylation profiles across the promoter CGIs of X-linked cat *KDM6A*, *MED14*, and *DDX3X*, which are the known XCI escapers[39]. The CGIs are highlighted by blue boxes. The WGBS data were from skin regions (cat #01) or peripheral blood cells (cats #04 and #10). The WGBS data of peripheral blood cells from a male cat named Boris was obtained from a publicly available dataset (SRX548709).

**Extended Data Figure 6. X-linked *Sly* of golden hamsters is distinct from *Arhgap36*.**

The genomic position of *Arhgap36* on a high-quality sequence assembly of the golden hamster X chromosome[44] (red bar, top) and the genetic mapping result of *Sly* (ref. 43) (orange box, bottom) are shown. The positions of the genetic markers used to map *Sly* are shown on the genome sequence and the genetic map.

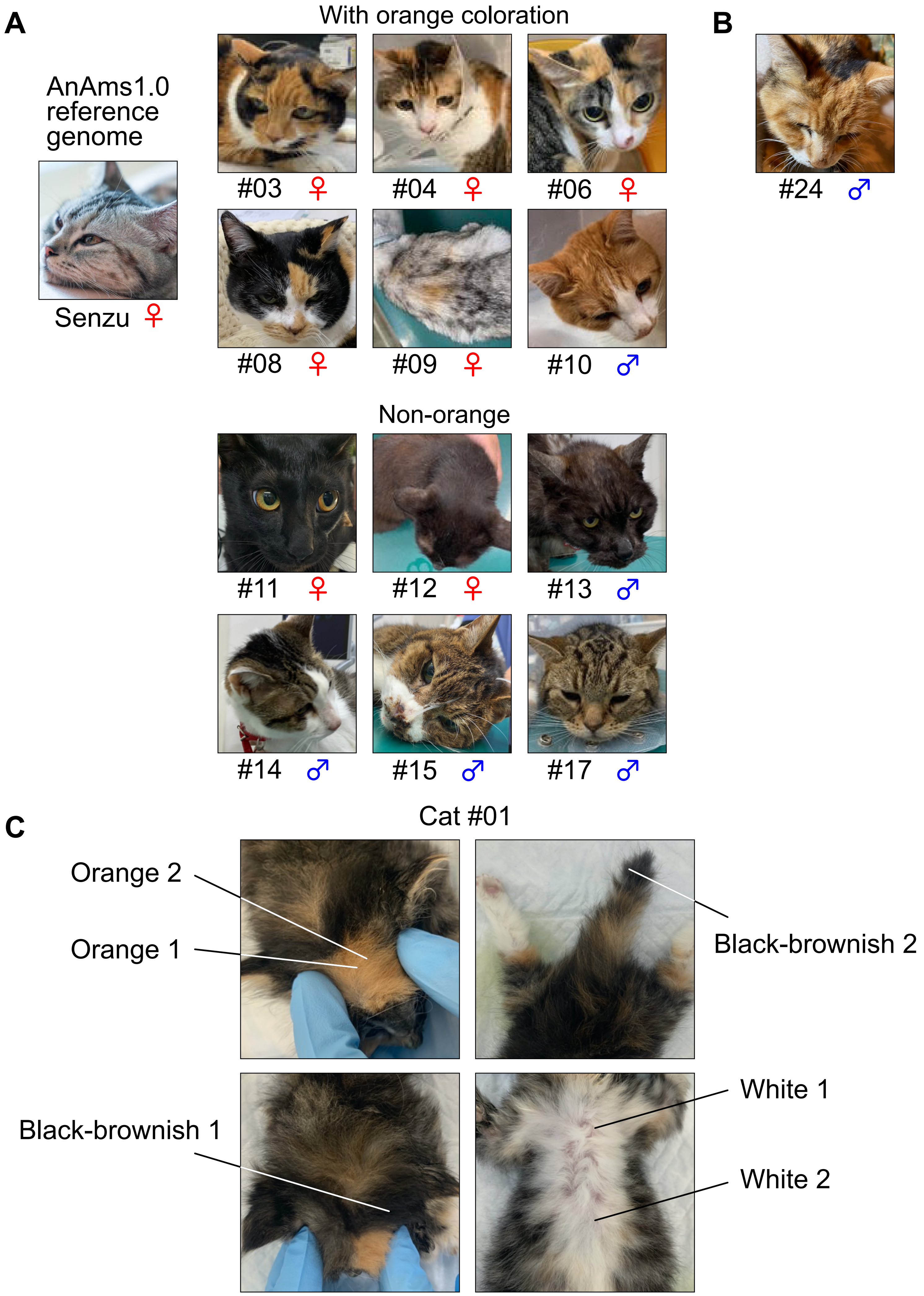


**Extended Data Figure 1**

**A**

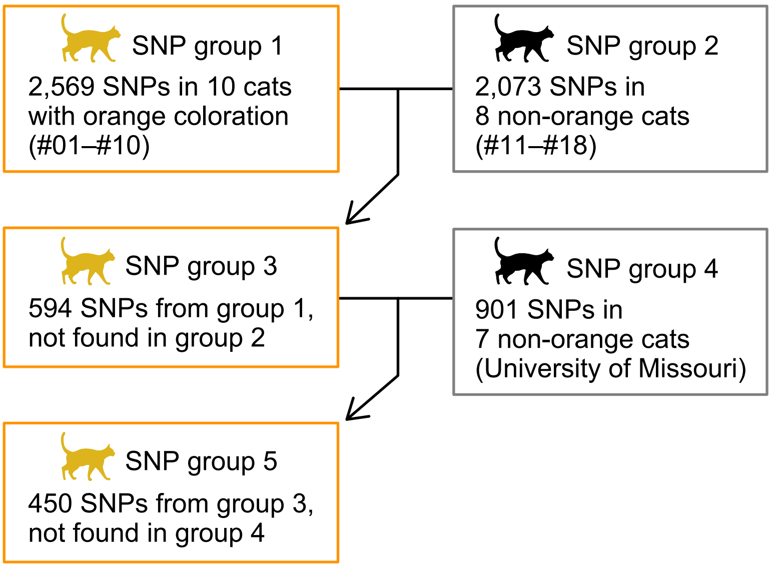


**B**

| | SNP group 1 | SNP group 3 | SNP group 5 |
|---|---|---|---|
| Frameshift variant | 2 | 0 | 0 |
| Missense variant | 4 | 0 | 0 |
| Nonsense variant | 0 | 0 | 0 |
| Synonymous variant | 5 | 1 | 1 |
| Intron variant | 634 | 183 | 136 |
| 5' prime UTR variant | 2 | 0 | 0 |
| 3' prime UTR variant | 9 | 3 | 3 |
| Intergenic region | 1,913 | 407 | 310 |
| Total | 2,569 | 594 | 450 |

**C**

| | SNP group 5 | *ZNF280C* | *SLC25A14* | *GPR119* | AnAmsX_07050 | *ENOX2* | *ARHGAP36* | *IGSF1* | *STK26* |
|---|---|---|---|---|---|---|---|---|---|
| Frameshift variant | 0 | 0 | 0 | 0 | 0 | 0 | 0 | 0 | 0 |
| Missense variant | 0 | 0 | 0 | 0 | 0 | 0 | 0 | 0 | 0 |
| Nonsense variant | 0 | 0 | 0 | 0 | 0 | 0 | 0 | 0 | 0 |
| Synonymous variant | 1 | 0 | 0 | 0 | 0 | 1 | 0 | 0 | 0 |
| Intron variant | 136 | 9 | 11 | 3 | 1 | 74 | 24 | 2 | 12 |
| 5' prime UTR variant | 0 | 0 | 0 | 0 | 0 | 0 | 0 | 0 | 0 |
| 3' prime UTR variant | 3 | 0 | 0 | 0 | 2 | 0 | 0 | 1 | 0 |
| Intergenic region | 310 | - | - | - | - | - | - | - | - |
| Total | 450 | | | | | | | | |

| Cat ID | *ZNF280C* | *SLC25A14* | *GPR119* | AnAmsX_07050 | *ENOX2* | *ARHGAP36* | *IGSF1* | *STK26* |
|---|---|---|---|---|---|---|---|---|
| #01 | 0 | 0 | 0 | 0 | 5 | 4 | 1 | 8 |
| #02 | 0 | 0 | 0 | 0 | 4 | 6 | 1 | 3 |
| #03 | 1 | 1 | 0 | 0 | 4 | 3 | 1 | 1 |
| #04 | 3 | 6 | 3 | 2 | 48 | 14 | 0 | 3 |
| #05 | 0 | 2 | 0 | 1 | 1 | 1 | 1 | 3 |
| #06 | 2 | 3 | 0 | 0 | 4 | 1 | 0 | 0 |
| #07 | 3 | 0 | 0 | 0 | 6 | 3 | 0 | 0 |
| #08 | 0 | 0 | 0 | 0 | 1 | 2 | 0 | 1 |
| #09 | 4 | 5 | 3 | 2 | 51 | 14 | 0 | 2 |
| #10 | 3 | 0 | 0 | 0 | 0 | 1 | 0 | 1 |

**D**

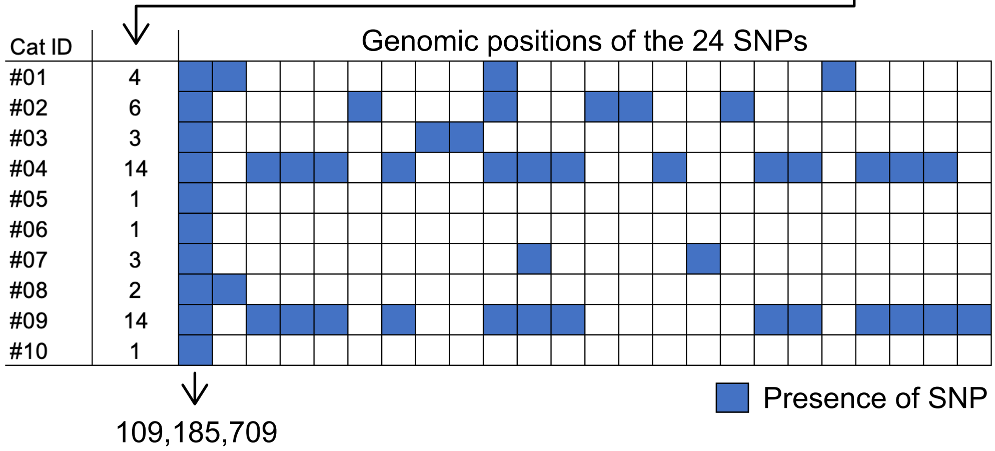


**Extended Data Figure 2**

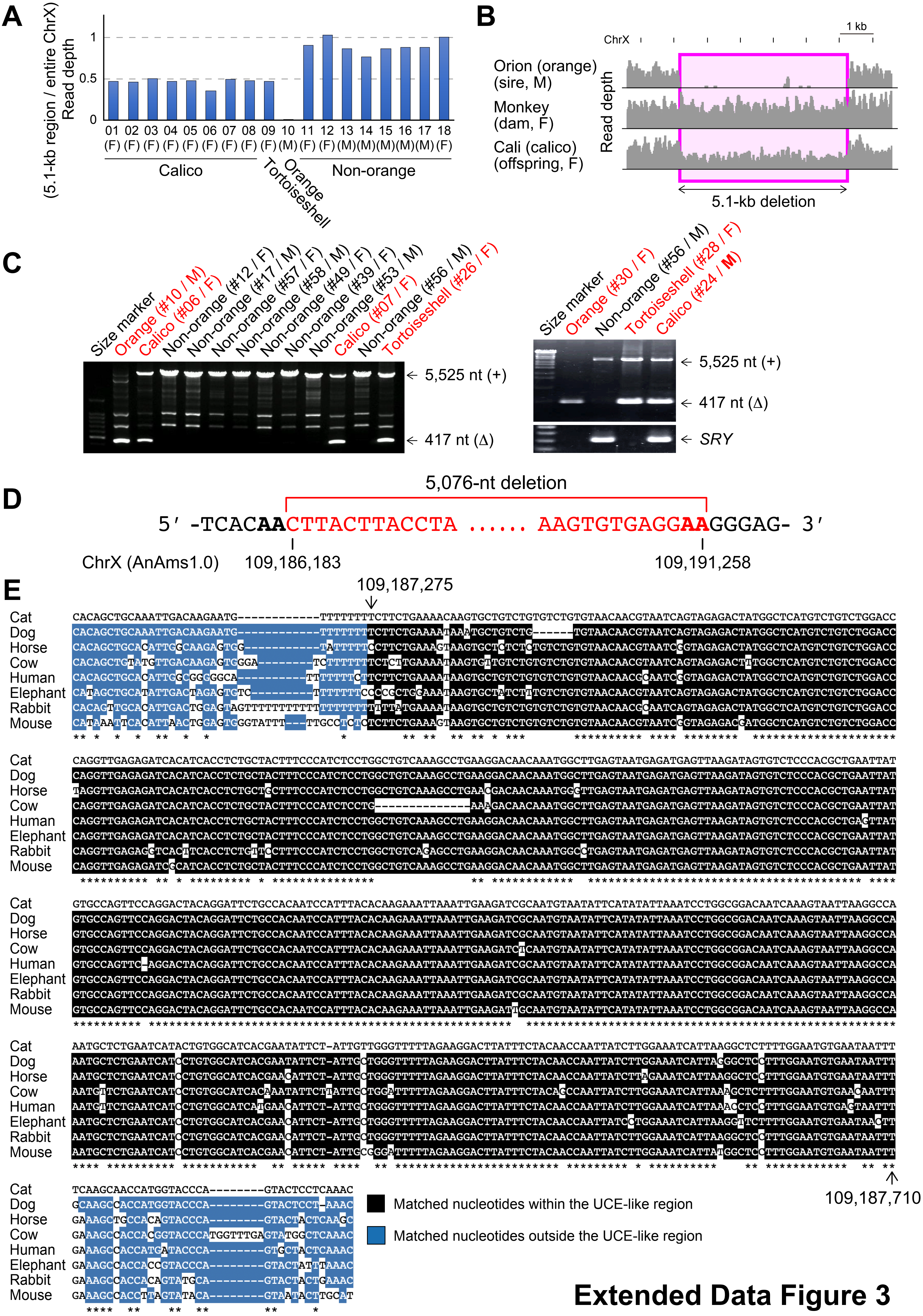


Extended Data Figure 3

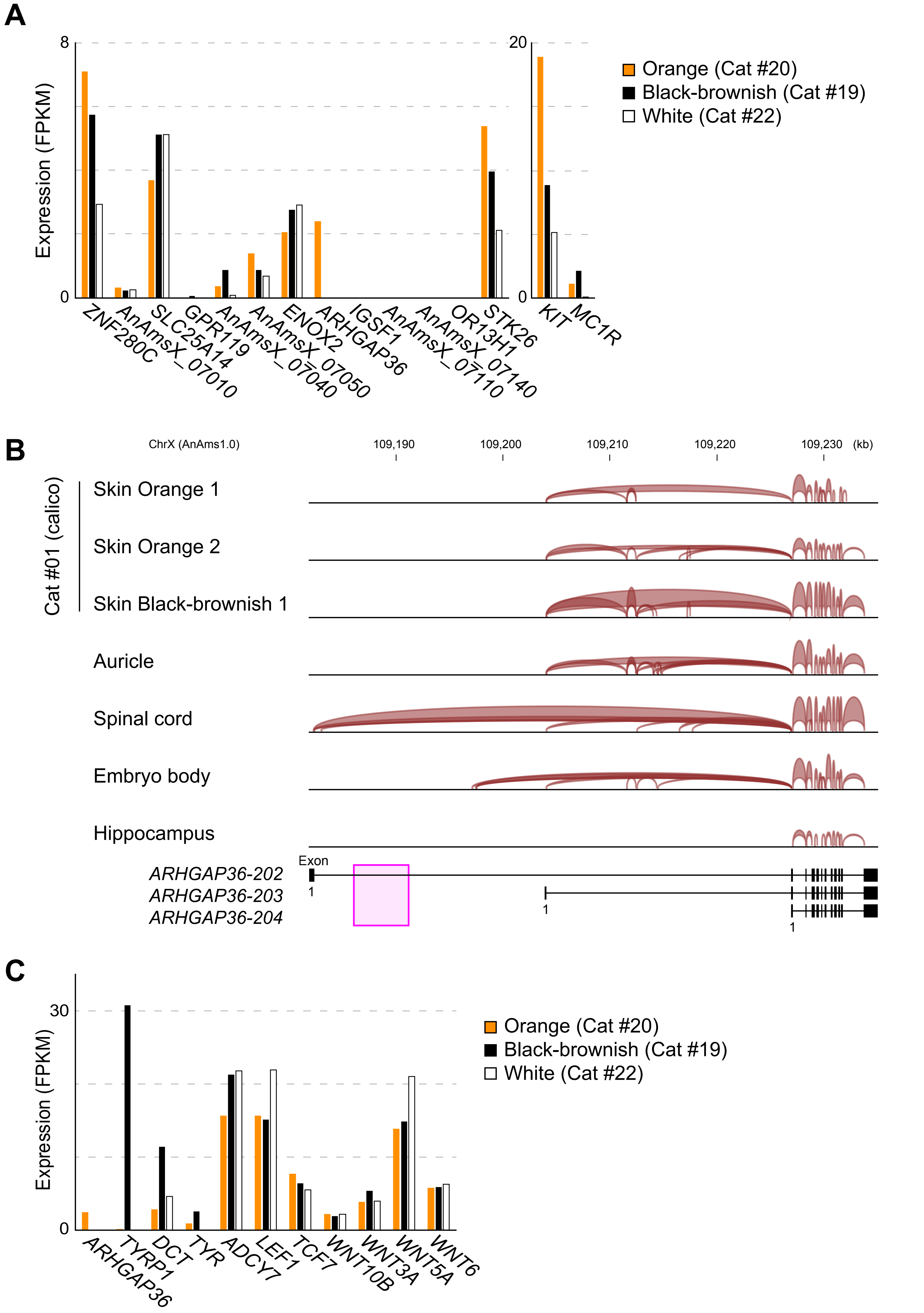


**Extended Data Figure 4**

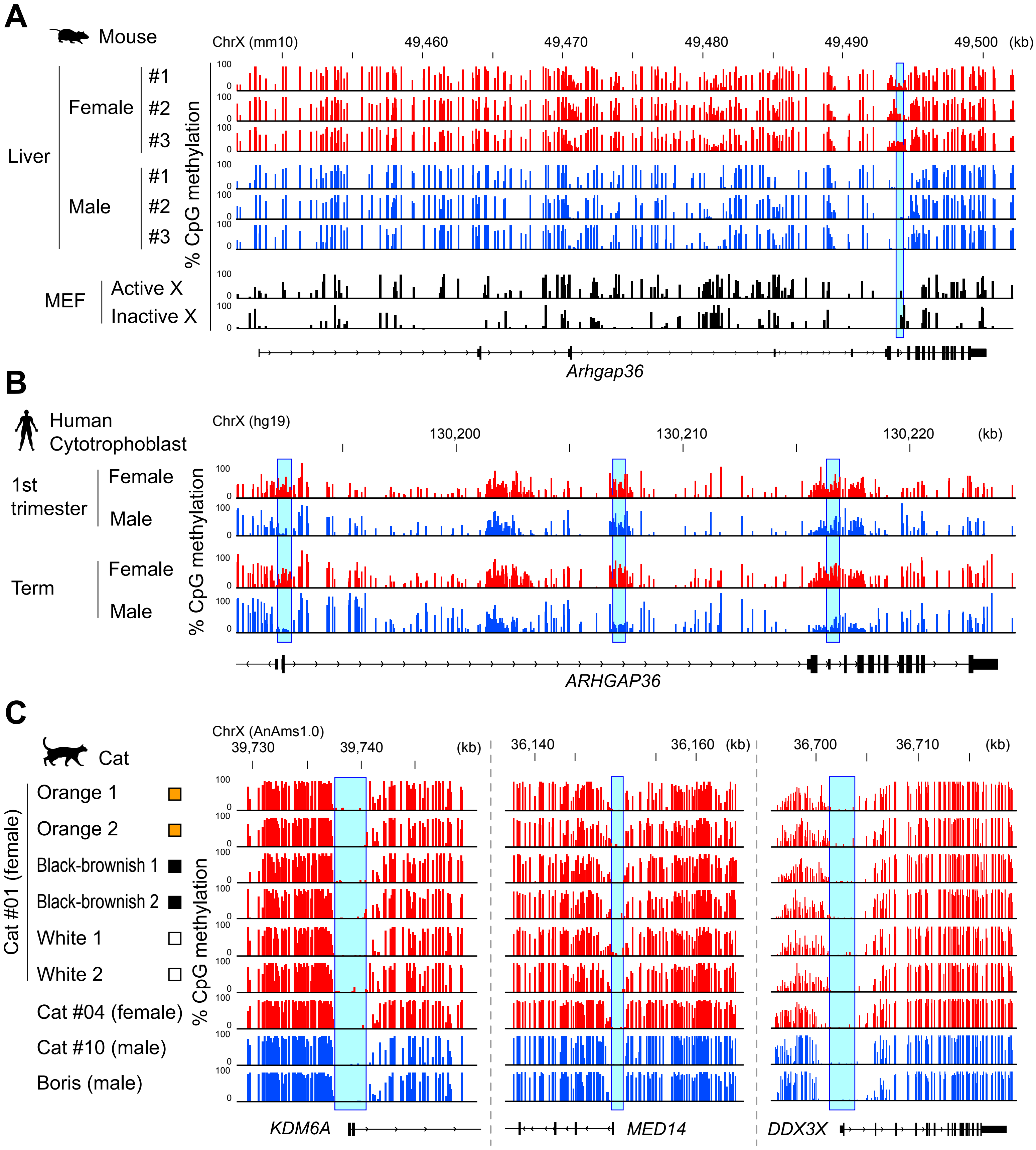


**Extended Data Figure 5**

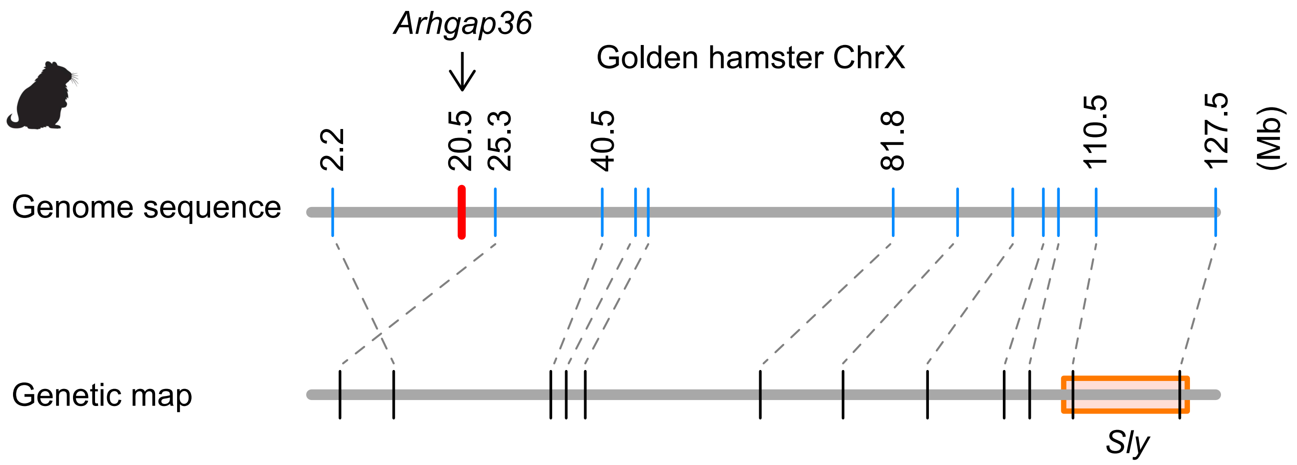


**Extended Data Figure 6**
